# Association of BDNF Val66Met Polymorphism and Brain BDNF levels with Major Depression and Suicide

**DOI:** 10.1101/233304

**Authors:** Mariam M. Youssef, Mark D. Underwood, Yung-Yu Huang, Shu-chi Hsiung, Yan Liu, Norman R. Simpson, Mihran J. Bakalian, Gorazd B. Rosoklija, Andrew J. Dwork, Victoria Arango, J. John Mann

## Abstract

Brain-derived neurotrophic factor (BDNF) is implicated in the pathophysiology of major depressive disorder (MDD) and suicide. Both are partly caused by early life adversity (ELA) and ELA reduces both BDNF protein and gene expression. This study examines the association of BDNF Val66Met polymorphism and brain BDNF levels with depression and suicide. We hypothesized that both MDD and ELA would be associated with the Met allele and lower brain BDNF levels. Such an association would be consistent with low BDNF mediating the effect of ELA on adulthood suicide and MDD. BDNF Val66Met polymorphism was genotyped in postmortem brains of 37 suicide decedents and 53 non-suicides. Additionally, BDNF protein levels were determined by Western blot in dorsolateral prefrontal cortex (Brodmann area 9; BA9), anterior cingulate cortex (ACC; BA24), caudal brainstem and rostral brainstem. The relationships between these measures and major depression, death by suicide and reported childhood adversity were examined. Depressed subjects had an excess of the Met allele and lower BDNF levels in ACC and caudal brainstem compared with non-depressed subjects. No effect of history of suicide death or early life adversity was observed with genotype, but lower BDNF levels in ACC were found in subjects who had been exposed to early life adversity and/or died by suicide compared to nonsuicide decedents and no reported childhood adversity. This study provides further evidence for low BDNF in major depression related to the BDNF met risk allele. Future studies should seek to determine how altered BDNF expression contributes to MDD and suicide.

## 1. Introduction

Suicide accounted for almost 1 million deaths worldwide in 2015 (WHO, 2016). Major Depressive Disorder (MDD) is the most prevalent psychiatric disorder in suicide decedents (Cavanagh *et al*, 2003). The stress diathesis model of suicidal behavior (Mann, 2003; Mann *et al*, 1999; van Heeringen and Mann, 2014) posits that the risk for suicide is determined, not only by the underlying psychiatric illness (stressor), but also by a trait-like diathesis. Identifying possible biological causes for this diathesis may aid in the development of prediction, prevention and treatment strategies for suicide risk.

Some studies have reported thinner prefrontal and anterior cingulate cortex in depressed suicide attempters (Wagner *et al*, 2011; Wagner *et al*, 2012), lower density of neurons in dorsal and ventral prefrontal cortex (Underwood *et al*, 2012) and fewer mature granule cells in the dentate gyrus of depressed suicides (Boldrini *et al*, 2013), raising the possibility of a deficit in brain neurotrophic pathways. Brain-derived neurotrophic factor (BDNF) is one of the neurotrophins (Huang and Reichardt, 2001) which regulate neuron survival, plasticity (Berton *et al*, 2006; McAllister, 2001; Morse *et al*, 1993; Poo, 2001; Tsankova *et al*, 2006) and synaptic function (Lessmann et al., 2003). BDNF plays an integral role in differentiation during development (Alcantara *et al*, 2006; Engelhardt *et al*, 2007), is regulated by stress (Roceri *et al*, 2004) and is associated with the pathophysiology of mental disorders, particularly major depression (Russo-Neustadt, 2003). Thus, BDNF is a candidate molecule for contributing to the adverse brain effects of exposure to early life stress that may mediate the effects on adulthood risk of major depression and suicide.

A functional polymorphism (rs6265) in the BDNF gene results in a valine-to-methionine substitution at codon residue 66 (Val66Met) (Post, 2007; Rybakowski, 2008; Schumacher *et al*, 2005). The *Met* allele is associated with less BDNF activity (Egan *et al*, 2003) and lower serum levels (Ozan *et al*, 2010) and appears to be associated with major depression (Hwang *et al*, 2006), memory impairments (Egan *et al*, 2003; Hariri *et al*, 2003), reduced hippocampal activity (Chen *et al*, 2004) and anxiety-related behaviors in animal models (Chen *et al*, 2006), although not all studies agree (Chen *et al*, 2008; Cohen *et al*, 2004; Schumacher *et al*, 2005; Sen *et al*, 2003; Strauss *et al*, 2005; Surtees *et al*, 2007). Furthermore, an excess of the *Met* allele increases the risk for suicidal behavior (Iga *et al*, 2007; Schenkel *et al*, 2010), particularly in depressed patients (Sarchiapone *et al*, 2008) and those exposed to early life stress (Pregelj *et al*, 2011).

Low plasma BDNF levels are reported in major depression (de Azevedo Cardoso *et al*, 2014; Karege *et al*, 2002; Sen *et al*, 2008) and suicidal behavior (Kim *et al*, 2007). Postmortem studies indicate low BDNF protein levels in amygdala (Guilloux *et al*, 2012) and decreased BDNF signaling in anterior cingulate cortex (Tripp *et al*, 2012) of depressed patients. Low BDNF protein was also reported in hippocampus (Banerjee *et al*, 2013; Dwivedi *et al*, 2003; Karege *et al*, 2005) and prefrontal cortex (Dwivedi *et al*, 2003; Karege *et al*, 2005) of suicide decedents. Antidepressants increase BDNF blood levels in depressed patients (Chen *et al*, 2001; Gervasoni *et al*, 2005; Gonul *et al*, 2005) and animal models of depression (Nibuya *et al*, 1995; Russo-Neustadt *et al*, 1999). Moreover, in animal depression models, injection of BDNF into the hippocampus (Shirayama *et al*, 2002) and midbrain (Siuciak *et al*, 1997) produces an antidepressant-like effect (e.g. decreased escape failure in the learned helplessness paradigm and decreased immobility in the forced swim test).

Exposure to stress decreases brain BDNF levels in rodents (Nibuya *et al*, 1999; Roceri *et al*, 2004; Smith *et al*, 1995). Stress, particularly in early life, can down-regulate BDNF in depressed (Grassi-Oliveira *et al*, 2008) and suicidal patients (Dwivedi, 2010). Some, but not all (Perroud *et al*, 2008) studies, show that exposure to early life stress in BDNF Met carriers predicts future depression (Aguilera *et al*, 2009; Gatt *et al*, 2009) and suicide (Pregelj *et al*, 2011).

In the current study, we sought to identify the inter-relationship of BDNF in suicide, major depression and reported childhood adversity by examining the BDNF polymorphism and BDNF protein levels in prefrontal cortex (PFC; Brodmann Area 9), anterior cingulate cortex (ACC; Brodmann area 24) and brainstem *postmortem* in cases of major depression, suicide decedents and nonsuicide, nopsychiatric sudden death comparison groups. We hypothesized that major depression would be associated with the Met allele and lower brain BDNF levels and this association would be more pronounced in suicide and/or MDD. We also hypothesized that reported childhood adversity would also correlate with low BDNF protein level in depressed suicide decedents, consistent with a model that posits low BDNF expression at least partly mediates the effect of early life adversity on adult risk of major depression and suicide.

## 2. Methods

### 2.1. Subjects

Brain samples from 37 suicide decedents and 53 nonsuicide comparison subjects were studied and clinical and demographic details are in Table 1. Procedures for collection and use of brain tissue were approved by the applicable Institutional Review Boards.

**Table 1:** Demographic and Clinical Characteristics of non-MDD and MDD Subjects:

| Variable | Non-MDD<br>(N= 45) |  | MDD<br>(N= 45) |  | Analysis<br>*p* * |
| --- | --- | --- | --- | --- | --- |
|  | *Mean* | *SD* | *Mean* | *SD* |  |
| Age (years) | 40.4 | 17.7 | 47.1 | 17.8 | 0.077 |
| Alcohol level (%) | 0.0143 | 0.0545 | 0.0079 | 0.0269 | 0.520 |
| pH of Brain Tissue | 6.5 | 0.3 | 6.5 | 0.3 | 0.638 |
| Total Aggression Score | 13.1 | 5.4 | 13.7 | 4.3 | 0.576 |
| Postmortem Interval | 15.2 | 4.8 | 16.3 | 6.7 | 0.413 |
| RIN | 6.5 | 0.6 | 6 | 1 | 0.280 |
|  | *N* | *%* | *N* | *%* |  |
| Sex: |  |  |  |  |  |
| • Female | 6 | 13.3 | 10 | 22.2 | 0.409 |
| • Male | 39 | 86.7 | 35 | 77.8 |  |
| Race: |  |  |  |  |  |
| • White | 35 | 77.8 | 37 | 84.1 | 0.792 |
| • Black | 5 | 11.1 | 4 | 9.1 |  |
| • Hispanic | 5 | 11.1 | 3 | 6.8 |  |
| Tobacco Smoking: |  |  |  |  |  |
| • Yes | 18 | 43.9 | 19 | 57.6 | 0.350 |
| • No | 23 | 56.1 | 14 | 42.4 |  |
| Death by suicide |  |  |  |  |  |
| • Yes | 5 | 11.1 | 32 | 71.1 | < **0.001** |
| • No | 40 | 88.9 | 13 | 28.9 |  |
| Early Life Adversity |  |  |  |  |  |
| • Yes | 15 | 33.3 | 17 | 39.5 | 0.658 |
| • No | 30 | 66.7 | 26 | 60.5 |  |
| History of Personality Disorder |  |  |  |  |  |
| • Yes | 5 | 11.1 | 8 | 18.6 | 0.378 |
| • No | 40 | 88.9 | 35 | 81.4 |  |

\* Independent Sample T-test (df=81, df=88 for Age) for continuous variables; Chi-square analysis (df=1) for categorical variables. Abbreviations: MDD= major depressive disorder, RIN= RNA integrity number, SD= standard deviation.

All suicide and nonsuicide subjects died suddenly, without a prolonged agonal period that might have an impact on the biological measures. Postmortem interval (PMI, time from death to freezing of brain samples) was limited to 24 hours. Brain samples were coded and assayed by laboratory personnel blind to the cause of death. The brainstem was dissected from the forebrain, which was bisected and cut into 2 cm slabs with the tissue blocks frozen for later sectioning. The brainstem was cut in a cryostat in the horizontal plane into 20 μm thick sections. Sections were selected that contained rostral and caudal levels of the dorsal raphe nucleus, the brain region containing serotonerigic neurons that innervate the forebrain. The dorsal raphe nucleus was selected as a region of interest for the actions of BDNF because of the role of serotonin and serotonin-synthesizing neurons in mood disorders and suicide, and the reported relationship between BDNF and serotonergic neurons (Martinowich and Lu, 2008). The rostral and caudal levels are referred to as rostral and caudal brainstem, respectively. Toxicological screening of body fluids (blood, bile, aqueous humor, and urine) as well as brain tissue was performed for cocaine, opiates, alcohol, cannabinoids, and other acidic and basic drugs, and major psychotropic drugs.

Psychological autopsies were completed and psychiatric diagnoses made using our validated method as previously described (Kelly and Mann, 1996). In brief, after giving written informed consent, at least one next of kin of subjects was interviewed. The Structured Clinical Interview for DSM-IV Axis I (SCID-I) (First *et al*, 1995), Structured Clinical Interview for DSM-IV Axis II Personality Disorders (SCID-II) (First *et al*, 1997), Brown-Goodwin Lifetime History of Aggression Scale (Brown *et al*, 1979) and Columbia Suicide History Form (Oquendo *et al*, 2003) were administered to the informant by a psychologist with at least a Master’s level degree. The Brown-Goodwin Lifetime History of Aggression Scale score was calculated, excluding the item regarding self-inflicted injury. Research diagnoses were based on DSM-IV criteria. The sample was divided clinically into subjects with a history of DSM-IV major depressive disorder (MDD) (N=45) and those without such history (non-MDD) (N=45). History of early life adversity in terms of physical or sexual abuse or neglect before the age of 15 years was also obtained during the psychological autopsy interview using a checklist.

Exclusion criteria included: 1) cases of uncertain manner of death, 2) gross neuropathology, 3) positive toxicology screens for psychoactive and neurotoxic drugs, 4) alcohol use disorder and 5) history of bipolar disorder or psychosis.

### 2.2. Genotyping

DNA was extracted from frozen brain tissue. The BDNF Val66Met polymorphism (GenBank dbSNP: rs6265) was typed by PCR protocol as published previously (Sublette *et al*, 2008). Briefly, the oligonucleotide primers, sense MannBF-1F (5’-ATCCCGGTGAAAGAAAGCCCTAAC-3’) and antisense MannBF-1R (5’-CCCCTGCAGCCTTCTTTTGTGTAA-3’), were used to amplify a PCR fragment of 673 bp length. PCR was carried out in a 20 μl volume, containing 100 ng DNA, 40 ng of each primer with HotStartTaq Plus Master Mix kit (Qiagen). Samples were processed in a BioRad T100 Thermal Cycler (BioRad, USA). DNA samples were denatured first at 95°C for 6 min. Thirty temperature cycles, consisting of 30 sec at 95°C, 40 sec at 60°C, and 40 sec at 72°C, were followed by a final extension step of 72°C for 4 min. The PCR fragments were digested with *BsaA I* restriction enzyme (NE Biolab, MA, USA), which produces 3 fragments of 275, 321 and 77 bp when guanine is present at nucleotide 1249, and 2 fragments of 321 and 352 bp if cytosine is present at this position. The digested PCR products were separated on a 1.2% agarose gel.

### 2.3. Western blotting of BDNF

Frozen brain samples from Brodmann Area (BA) 9 of 85 subjects, BA24 of 51 subjects, caudal brainstem of 33 subjects and rostral brainstem of 25 subjects were homogenized in cell lysis buffer (Cell Signaling, Beverly, MA, USA) plus 2% phosphatase inhibitor cocktail 2, 2% phosphatase inhibitor cocktail 3 and 2% Protease Inhibitor cocktail (Sigma, St. Louis, MO, USA). The homogenate was centrifuged for 10 min at 15,000 x g at 4°C and the supernatant was transferred into Eppendorf tubes and stored at −80°C. Protein concentration was determined using Beckman DU530 spectrophotometer. Laemmli buffer with β-mercaptoethanol was added in Eppendorf tubes. After adding protein samples, water was added up to 20 μl to each tube. The samples were denatured for five minutes at 95°C and then centrifuged for five seconds. Protein samples (25 μg) were loaded onto 12% Mini-PROTEAN TGX gel. The gel was run with 10x tris-glycine with SDS for 20 minutes at 40 volts and increased to 100 volts for 1 hour and transferred in an enhanced chemiluminescence (ECL) nitrocellulose membrane (Amersham, Arlington Heights, IL, USA) for 1 hour and 15 minutes at 90 volts with a transfer buffer (1x tris-glycine with 20% methanol) at 4°C. The membranes were washed with TBST buffer [10 mm Tris-base, 0.15 m NaCl, and 0.1% Tween-20] for 10 min. The blots were blocked by incubating with 5% non-fat milk in TBST for 1 hour. Then the blots were incubated overnight at 4°C with primary polyclonal anti-BDNF antibody (Alomone Labs, Jerusalem, Israel) at a dilution of 1:50,000. The membranes were washed with TBST and incubated with horseradish-peroxidase secondary antibody [anti-rabbit immunoglobulin G (IgG); 1:5000] in 5% non-fat milk for 3 h at 4°C. The membranes were extensively washed with TBST and exposed to ECL autoradiography film. The same nitrocellulose membrane was stripped and re-probed with GAPDH antibody (Cell signaling, Beverly, MA, USA).

### 2.4. Statistical analysis

The genotype frequency distributions of the SNPs were tested for Hardy–Weinberg equilibrium (HWE) in the non-MDD, nonsuicide controls.

Data were statistically evaluated using IBM SPSS Statistics (SPSS Inc., Chicago, Illinois, USA, Version 23.0). Comparison of demographic, clinical and postmortem characteristics between the depressed and non-depressed subjects were performed using two-tailed *t*-tests for continuous variables and Chi-square analyses for categorical variables. The differences in allele and genotype frequency distributions between the non-MDD *vs*. MDD, nonsuicides *vs*. suicides, as well as subjects with no history of early life adversity *vs*. those with such history were calculated using Pearson’s Chi-square tests (df = 1 and df = 2, respectively). The difference in life time history of aggression among the 3 genotype groups was assessed using one-way analysis of variance (ANOVA).

For BDNF protein analyses, we first explored the effects of age, sex, RNA integrity number (RIN), PMI and brain pH on the levels of BDNF protein using Pearson’s correlation analyses. No effect of these variables was detected on BDNF protein levels in the four brain regions studied, therefore, they were not added as covariates in further analyses. A 2-tailed t-test was used to compare BDNF protein levels between subjects with GG genotypes *vs*. carriers of A allele (AA and AG), and non-MDD *vs*. MDD in the four brain regions. We found an effect of depression diagnosis on BDNF protein levels, so we controlled for depression status in subsequent analyses (suicide *vs*. non-suicide, history of early life adversity *vs*. no early life adversity) using an analysis of covariance (ANCOVA) using MDD diagnosis as a co-variate. Additionally, we further classified the sample into four groups based on the manner of death (suicide or not) and history of exposure to early life adversity (non-suicide, no adversity *vs*. nonsuicide, adversity *vs*. suicide, no adversity *vs*. suicide, adversity) and examined the differences of BDNF protein levels among these four groups while controlling for depression status using ANCOVA in each region separately. Correction for multiple comparisons was made using the Bonferoni method.

## 3. Results

### 3.1. Demographics and Clinical Characteristics

Demographic and clinical characteristics of the study sample (N= 90) are shown in *Table 1*. The MDD and non-MDD groups did not differ in terms of age, sex, or race. The MDD group were more likely to die by suicide than the non-MDD group (χ^2^= 33.46, df= 1, p< 0.001). There were also no group differences in blood alcohol level, brain tissue pH, PMI and RIN. There were also no group differences in lifetime history of aggression score, reported early life adversity, or comorbid personality disorders.

### 3.2. BDNF Val66Met polymorphism allele and genotype frequencies

Genotype frequencies in the non-MDD, nonsuicide control group were in Hardy–Weinberg equilibrium (χ^2^ = 0.1117). BDNF Val66Met genotype distribution differed between MDD and non-MDD groups (χ^2^= 7.91, df= 2, p= 0.019), and post hoc testing showed that this was due to an excess of the Met allele (A) in the MDD group (χ2= 6.54, df= 1, p= 0.011) (see *Table 2*). The BDNF polymorphism did not differ between suicide and non-suicide decedents (χ2= 2.8, df= 2, p= 0.24, *Table 2*). Reported early life adversity was not associated with genotype (χ2= 0.96, df= 2, p= 0.62, *Table 2*). Sub-classifying the carriers of Met allele into two groups according to reported exposure to early life adversity revealed no association with major depression (χ2= 0.09, df= 1, p= 0.77, *Table 3*). The lifetime history of aggression score was not different between the three genotypes (F= 1.42, df= 2,79, p= 0.25).

**Table 2:** BDNF Val66Met polymorphism allele and genotype frequencies:

| rs6265 | N | Genotype N (%) | | | $\chi^2$* | P | Allele N (%) | | $\chi^2$* | P |
| --- | --- | --- | --- | --- | --- | --- | --- | --- | --- | --- |
|  |  | GG | AG | AA |  |  | G | A |  |  |
| **Non-MDD** | 45 | 34<br>(75.6) | 10<br>(22.2) | 1<br>(2.2) | 7.91 | **0.019** | 78<br>(87) | 12<br>(13) | 6.54 | **0.011** |
| **MDD** | 45 | 21<br>(46.7) | 22<br>(48.9) | 2<br>(4.4) |  |  | 64<br>(71) | 26<br>(29) |  |  |
| **Non-Suicide** | 53 | 36<br>(67.9) | 16<br>(28.3) | 1<br>(1.8) | 2.8 | 0.24 | 88<br>(83) | 18<br>(17) | 2.64 | 0.1 |
| **Suicide** | 37 | 19<br>(51.4) | 16<br>(43.2) | 2<br>(5.4) |  |  | 54<br>(73) | 20<br>(27) |  |  |
| **No early life adversity** | 58 | 37<br>(63.8) | 19<br>(32.8) | 2<br>(3.4) | 0.96 | 0.62 | 93<br>(80) | 23<br>(20) | 0.32 | 0.57 |
| **Early life adversity** | 32 | 18<br>(56.3) | 13<br>(40.6) | 1<br>(3.1) |  |  | 49<br>(77) | 15<br>(23) |  |  |

\*Chi-square analysis (df=1 for allele frequency comparisons, df= 2 for genotype frequency comparisons). Abbreviations: MDD= major depressive disorder.

**Table 3:** Distribution of BDND Met Carriers with and without History of Early Life Adversity in Relation to MDD:

| **Met Carriers** | **Non- MDD**<br>**(N = 11)** | **MDD**<br>**(N = 24)** | **Analysis**<br>***p*\*** |
| --- | --- | --- | --- |
| **With ELA** | 4 (36.4%) | 10 (41.7%) | 0.77 |
| **Without ELA** | 7 (63.6%) | 14 (58.3%) |  |

\*Chi-square analysis (df = 1). Abbreviations: MDD= major depressive disorder, ELA= early life adversity.

### 3.3. BDNF Protein

No statistically significant effects of age, sex, RIN, PMI, or brain pH on the levels of BDNF protein were detected (p > 0.05 all variables) (see supplementary data and *supplementary table*).

Subjects with GG genotype did not differ from carriers of A allele regarding BDNF levels in dlPFC (*t*_(83)_= −0.32, p= 0.8), ACC (*t*_(49)_= −0.73, p= 0.46), BSr (*t*_(23)_)= 0.16, p= 0.87) or BSc (*t*_(31)_= −2.02, p= 0.052). Likewise, there was no association between genotype (GG vs. A carrier) and BDNF level in depression, suicide or reported childhood adversity; multivariate analysis, mixed model analysis and stepwise linear regression analysis all failed to detect an effect of genotype.

However, BDNF protein level appeared to be lower in ACC (*t*_(49)_= 2.14, p= 0.04 without Bonferoni correction) and caudal brainstem (*t*_(31)_= 2.37, p= 0.02 without Bonferoni) in the MDD group compared with non-MDD group. No effect of major depression diagnosis was found on BDNF protein levels in dlPFC (*t*_(83)_= −0.72, p= 0.47) or rostral brainstem (*t*_(23)_= 0.58, p= 0.57) [see *figure 1*].

**Figure 1:**
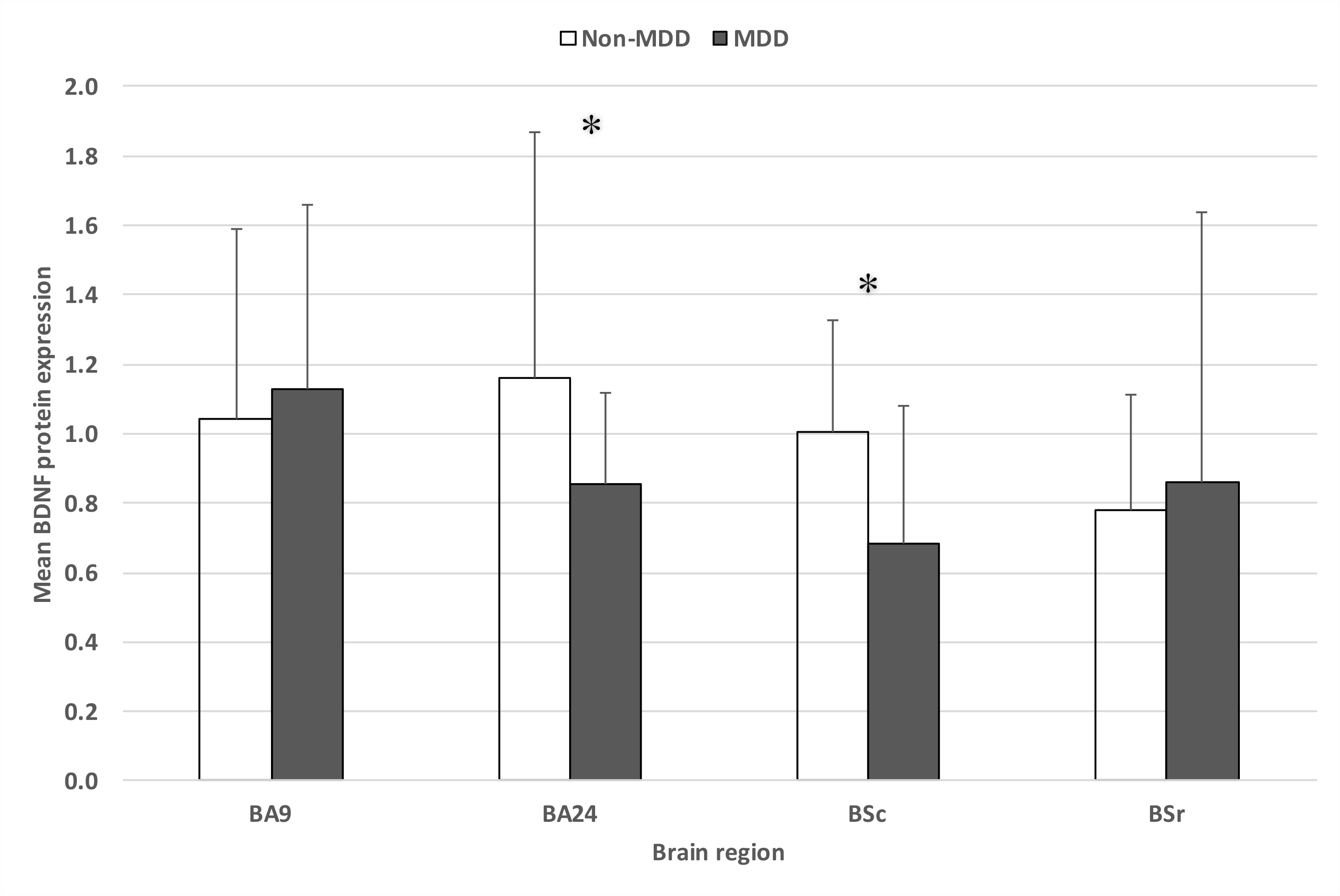
Bar chart of the mean BDNF protein expression in BA 9 (N= 85), BA 24 (N= 51), BSc (N= 33), and BSr (N= 25) in non-MDD and MDD groups. BDNF protein levels were lower in BA 24 and BSc of depressed patients compared to non-depressed subjects. No group differences were found in the BA 9 or in BSr. *p< 0.05. Abbreviations: BA= Brodmann area, BDNF= brain-derived neurotrophic factor, MDD= major depressive disorder, BSc= caudal brain stem, BSr= rostral brain stem.

Since there was an indication of a potential effect of MDD in ACC and caudal brainstem *(vide supra*), we controlled for MDD diagnosis in subsequent analyses of the effects of suicide and early life adversity by using ANCOVA. Classifying the study subjects based on manner of death (suicide or not) and exposure to early adversity or not (*figure 2*), BDNF protein level in ACC in suicide decendents with adversity history was lower compared with controls without reported adversity (*F*_(3, 46)_= 5.84, p= 0.006 uncorrected, p= 0.048 corrected), but not in the dlPFC (*F*_(3, 78)_= 0.89, p= 0.45), BSc (*F*_(2, 27)_= 0.48, p= 0.63), or BSr (*F*_(2, 20)_= 1.37, p= 0.28).

**Figure 2:**
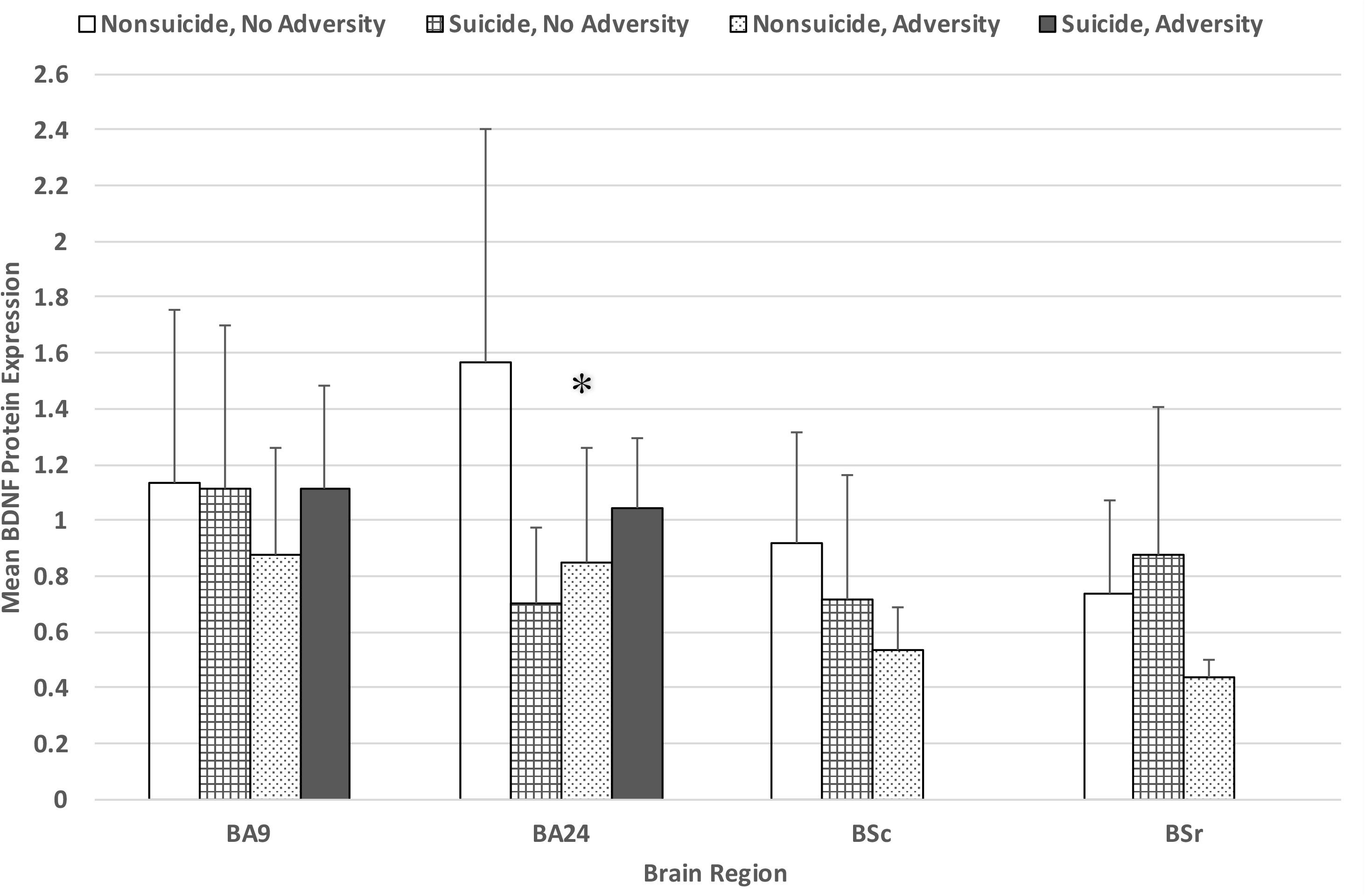
Bar chart of the mean BDNF protein levels in BA 9 (N= 85), BA 24 (N= 51), BSc (N= 33), and BSr (N= 25) in various groups. BDNF protein levels in ACC (BA24) in the group without history of suicide and adversity was higher than other groups. No group difference was found in the BA 9, BSc, or BSr. \**p*< 0.05. Abbreviations: BA= Brodmann area, BDNF= brain-derived neurotrophic factor, BSc= caudal brain stem, BSr= rostral brain stem.

There was no difference in BDNF levels between suicide decedents and nonsuicide decedents in dlPFC (*F*_(1,82)_= 0.004, p= 0.95), ACC (*F*_(1, 48)_= 0.52, p= 0.48), BSc (*F*_(1, 30)_= 0.09, p= 0.76) or BSr (*F*_(1, 22)_= 2.67, p= 0.12). Likewise, there was no difference in BDNF levels between subjects with reported early life adversity compared to those without such a history in dlPFC (*F*_(1, 79)_= 1.99, p= 0.16), ACC (*F*_(1, 48)_= 0.93, p= 0.34), BSc (*F*_(1, 28)_= 0.91, p= 0.35),or BSr (*F*_(1, 21)_= 1.89, p= 0.18).

## 4. Discussion

The current study provides evidence for a role of BDNF in the pathophysiology of MDD. We found an association of both BDNF Met allele and lower brain BDNF protein level in the ACC and caudal brainstem with MDD, and there were lower BDNF levels in association with a reported history of childhood adversity and suicide as cause of death. Contrary to our hypothesis, we did not find BDNF brain levels associated with genotype.

### 4.1. BDNF polymorphism

Our finding that the Met allele is associated with history of MDD is consistent with a reported association with geriatric depression (Hwang *et al*, 2006) and combined anxiety and depression diagnoses (Jiang *et al*, 2005; Taylor *et al*, 2007). A meta-analysis of Verhagen *et al* (2010) also reported such association with depression in men only. Chen *et al* (2006) found increased anxiety-related behaviors in BDNF^Met/Met^ mice indicating a causal relationship. The BDNF Met allele is associated with less hippocampal activity (Egan *et al*, 2003; Frey *et al*, 2007; Hariri *et al*, 2003) and smaller volume (Duman, 2002; Frodl *et al*, 2007; Gonul *et al*, 2011; Pezawas *et al*, 2004; Szeszko *et al*, 2005). On the other hand, many studies did not find such association (Gratacos *et al*, 2007; Hong *et al*, 2003; Surtees *et al*, 2007; Tsai *et al*, 2003). The differences in results from these studies may be due to differences in study design, sample characteristics (e.g. age groups, ethnicity or race) or because of the clinical and biological heterogeneity of major depression (Akiskal *et al*, 1981). Strauss *et al* (2005) found an association of Val allele with childhood mood disorders, whereas Hwang *et al* (2006) found that the Met allele is related to geriatric depression. Some studies conclude that a gene-gene or gene-environment interaction is required for this BDNF polymorphism to contribute to major depression (Belsky *et al*, 2009; Kaufman *et al*, 2006; Wichers *et al*, 2008). We found association of excess Met allele with MDD and a trend for lower BDNF protein in the brain in MDD, however, genotype appears to be unrelated to BDNF protein. We did not find a relationship between childhood adversity and genotype suggesting that exposure to adversity in early life did not drive the Met allele-depression association.

We did not find an association between the val66met BDNF polymorphism and death by suicide, consistent with previous studies on suicide decedents (Pregelj *et al*, 2011; Ratta-Apha *et al*, 2013; Zarrilli *et al*, 2009). The Met allele is reportedly associated with nonfatal suicide attempts (Sarchiapone *et al*, 2008). This may add further evidence to the hypothesis that fatal and nonfatal suicide attempts are only partially overlapping phenomena with different underlying genomic components (Mann *et al*, 2009).

### 4.2. BDNF level in brain

The MDD group had a trend for lower BDNF protein level in ACC and caudal brainstem, but not in dlPFC or rostral brainstem, compared with the non-MDD group. Our findings in ACC are consistent with Tripp *et al* (2012) who found that BDNF receptor signaling and BDNF-dependent gene expression are lower in ACC of depressed compared to non-depressed subjects. That same study reported no differences in BDNF mRNA levels, which suggests that differences in BDNF protein levels may be due to defective translation of BDNF, but not to gene transcription. Therefore, the functional effect of BDNF genotype appears linked to BDNF protein (Chen *et al*, 2004; Egan *et al*, 2003) and not mRNA levels.

The ACC is implicated in the pathophysiology of mood disorders (Drevets *et al*, 2008). Functional neuroimaging studies show that the ACC, particularly its rostral/subgenual part, has a role in emotion regulation (Davidson *et al*, 2002; Vogt *et al*, 1992). Volume reduction in this area is a consistent finding in mood disorders (Drevets *et al*, 1997). Low BDNF levels in ACC may, therefore, contribute to its functional and structural deficits in MDD.

Our finding of low BDNF brainstem levels in depressed subjects is consistent with Kozicz *et al* (2008) who reported low BDNF expression in midbrain of male depressed patients. BDNF is a neurotrophic factor for dopaminergic neurons of the substantia nigra (Hyman and Hofer, 1991) and has a powerful positive effect on the survival of human and rat mesencephalic dopaminergic neurons in culture (Zhou *et al*, 1994). Chronic infusion of BDNF into the rat midbrain increase the turnover of serotonin and levels of noradrenaline in many forebrain areas including the neocortex, basal ganglia and hippocampus (Altar *et al*, 1994; Siuciak *et al*, 1996) and infusion into periaqueductal gray and dorsal raphe or into the substantia nigra have antidepressant-like effects in several behavioral tasks (Siuciak *et al*, 1997).

We did not find an effect of suicide, by itself as a cause of death, on BDNF protein levels across the four brain regions examined. However, we did find that suicides, when considered with respect to exposure to early life adversity, did have lower BDNF level in ACC, but not in the other brain regions. Some previous studies found lower BDNF protein levels in the PFC of suicide decedents (Dwivedi *et al*, 2003; Karege *et al*, 2005; Pandey *et al*, 2010). However, the findings of these studies may be confounded by the underlying psychiatric diagnosis or exposure to early life adversity. Our current study included only subjects with history of MDD and we controlled for MDD diagnosis status. The sample in Pandey *et al* (2010) study was exclusively teenage suicide decedents with potentially different characteristics, risk factors and pathophysiology from adult suicide (Brent *et al*, 1993). Our findings indicate that low brain BDNF levels may contribute to the pathophysiology of MDD which in turn is a major risk factor for suicide.

The connection between low BDNF, suicide and early life adversity may come about by a dysregulated stress response system in which BDNF translation is altered by the chronic effects of stress resulting from maltreatment in early life, and also with the acute effects of stress accompanying suicide behavior. Infant maltreatment results in methylation of BDNF DNA resulting in reduced BDNF gene expression in the adult prefrontal cortex (Roth *et al*, 2009). Interestingly, BDNF promoter/exon IV, which plays as critical role in BDNF gene regulation (Dennis and Levitt, 2005), is frequently hypermethylated in the Wernicke area of the postmortem brain of suicide subjects (Keller *et al*, 2010). Additionally, overactive hypothalamic-pituitary-adrenal axis has been linked to death by suicide (Mann *et al*, 2006) and abnormal stress response may result in part from a loss of neuronal plasticity that could be relevant in suicidal behavior (Duman and Monteggia, 2006). Therefore, in addition to its involvement in depression, BDNF may play an important role in the neurobiology of suicidal behavior in the context of a psychosocial stress (Deveci *et al*, 2007). However, further investigation in a larger sample is required in order to draw definitive conclusions about associations with suicide independently of a diagnosis of MDD.

### 4.3. Genotype and BDNF level

We did not find association between the BDNF Val66Met polymorphism and brain BDNF protein levels, consistent with the Lee *et al* (2005) study of temporal cortex of Alzheimer’s patients. Our finding is also consistent with other (Duncan *et al*, 2009; Yoshimura *et al*, 2011; Zou *et al*, 2010), but not all (Ozan *et al*, 2010), studies that examined the effect of BDNF Val66Met polymorphism on blood BDNF levels. However, we found excess Met allele frequency as well as low ACC and brainstem BDNF levels in MDD. It has been reported that Met allele contributes to the defect in activity-dependent BDNF secretion; such that Met carriers have abnormal intracellular trafficking and packaging of pro-BDNF reducing the depolarization-dependent secretion of the mature peptide (Chen *et al*, 2004; Egan *et al*, 2003). This suggests that the relationship between BDNF Val66Met polymorphism and brain BDNF levels is complex. This polymorphism may be in linkage disequilibrium with another unidentified functional polymorphism. This explanation is supported by Bhang *et al* (2011) who reported that healthy volunteers who were homozygous for S at 5-HTTLPR and the Met allele of the BDNF Val66Met polymorphism displayed significantly lower serum BDNF levels. Future studies should examine the effect of combination of functional polymorphisms on brain BDNF levels.

### 4.4. Limitations

Despite t evidence found in our study for the association of BDNF with MDD, there is uncertainty about the mechanism through which BDNF contribute to depressive pathophysiology. Alleles at the Val66Met locus may also be in linkage disequilibrium with other risk alleles in different genes. Future studies should genomic abnormalities more broadly in relation to MDD and suicide in order to better understand the mechanism underlying the causal role of genetic and early childhood adversity experiences.

## Acknowledgements

Mariam Youssef performed the statistical analysis and conceptualized and wrote the manuscript. Yung Yu assisted with BDNF polymorphism genotyping. Norman Simpson performed the DNA extraction. Yan Liu and Shu-chi Hisung conducted the Western Blot experiments. Mihran Bakalian assisted with the data analysis. Mark Underwood assisted with the data analysis and manuscript preparation. Gorazd Rosoklija and Andrew Dwork contributed to the brain collection, psychological autopsy and neuropathology assessment. Victoria Arango designed the project, assisted in data interpretation and manuscript editing. J John Mann designed the project, assisted in data interpretation and manuscript editing.

